# Loss of *PAX4* results in disrupted endocrine pancreas development and neonatal diabetes in pigs

**DOI:** 10.64898/2026.05.21.727014

**Authors:** Ravikanthreddy Poonooru, Ki-Eun Park, Amanda Schmelzle, Bhanu P. Telugu

## Abstract

Variants in the human *PAX4* gene are associated with both monogenic and complex forms of diabetes, yet their pathogenic effects remain difficult to define in models that accurately mimic human islet architecture and neonatal metabolic transitions. Here, we created a porcine *PAX4* loss-of-function model using CRISPR/Cas9 cytidine deaminase base editing to introduce a premature stop codon in the *PAX4* coding sequence. *PAX4* knockout piglets developed severe hyperglycemia within 24 hours of birth, followed by rapid postnatal clinical deterioration and uniform death by day 3. Biochemical analysis showed significant diabetic decompensation, including electrolyte imbalances, hyperosmolality, azotemia, dyslipidemia, and metabolic acidosis. Gross and histological examinations revealed notable pancreatic hypoplasia with preservation of exocrine tissue. Single-nucleus RNA sequencing and immunohistochemistry demonstrated an almost complete loss of insulin-and somatostatin-producing β-and δ-cells, respectively, with relative preservation of glucagon-expressing α-cells. Overall, these results establish *PAX4* as a crucial factor in pancreatic endocrine development and postnatal glucose regulation in a large-animal model. This platform offers a human-relevant system for studying diabetes-associated *PAX4* variants and for testing regenerative and gene-based therapies for insulin-deficient diabetes.

## Introduction

Diabetes mellitus is a “new age” epidemic characterized by a persistent increase in the circulating blood glucose levels (hyperglycemia) due to the loss or reduction in the number of functional primary beta (β) cells (*type 1 diabetes*), or reduced production/loss of responsiveness to insulin (*type 2 diabetes*) (1). Globally, diabetes is responsible for 4.6 million deaths per year. Nearly 600 million adults worldwide currently live with diabetes, a figure projected to surpass 850 million by 2050 (2). This trend highlights the urgent need to elucidate the developmental and genetic mechanisms governing pancreatic endocrine cell formation and maintenance. Dysfunction of insulin-producing β-cells is central to both monogenic and complex forms of diabetes, positioning the study of endocrine lineage specification as a primary objective in diabetes research and regenerative medicine (3).

The development of pancreatic endocrine cells is governed by a hierarchical transcriptional network that directs multipotent progenitors toward specific hormone-producing lineages. This framework has been largely defined through genetic studies in mice, where endocrine differentiation begins with the transient expression of the basic helix-loop-helix transcription factor Neurogenin 3 (*Ngn3*) (4). *Ngn3*-positive progenitors generate all endocrine cell types, including α-, β-, δ-, PP-, and ε-cells, by activating downstream transcription factors such as *NeuroD1*, *Nkx2.2*, *Rfx6*, and *Pax* genes, respectively (Fig.1A). Among the key regulatory genes, *PAX4* is a key determinant of endocrine cell fate decisions. *PAX4* encodes a paired-homeodomain transcription factor that directs endocrine progenitors toward β-and δ-cell lineages while repressing α-cell identity (5,6). In murine models, Pax4 deficiency leads to severe neonatal hyperglycemia, an almost complete absence of β-cells, and a corresponding increase in α-cells, underscoring its critical role in β-cell specification (7). In humans, genetic variation in *PAX4* is associated with both monogenic diabetes and increased risk of *type 2 diabetes*, including population-enriched missense variants linked to impaired β-cell function (3, 8). However, the developmental and physiological consequences of *PAX4* loss have not been investigated in a large-animal model that more accurately reflects human islet architecture and metabolic physiology.

**Figure 1.**
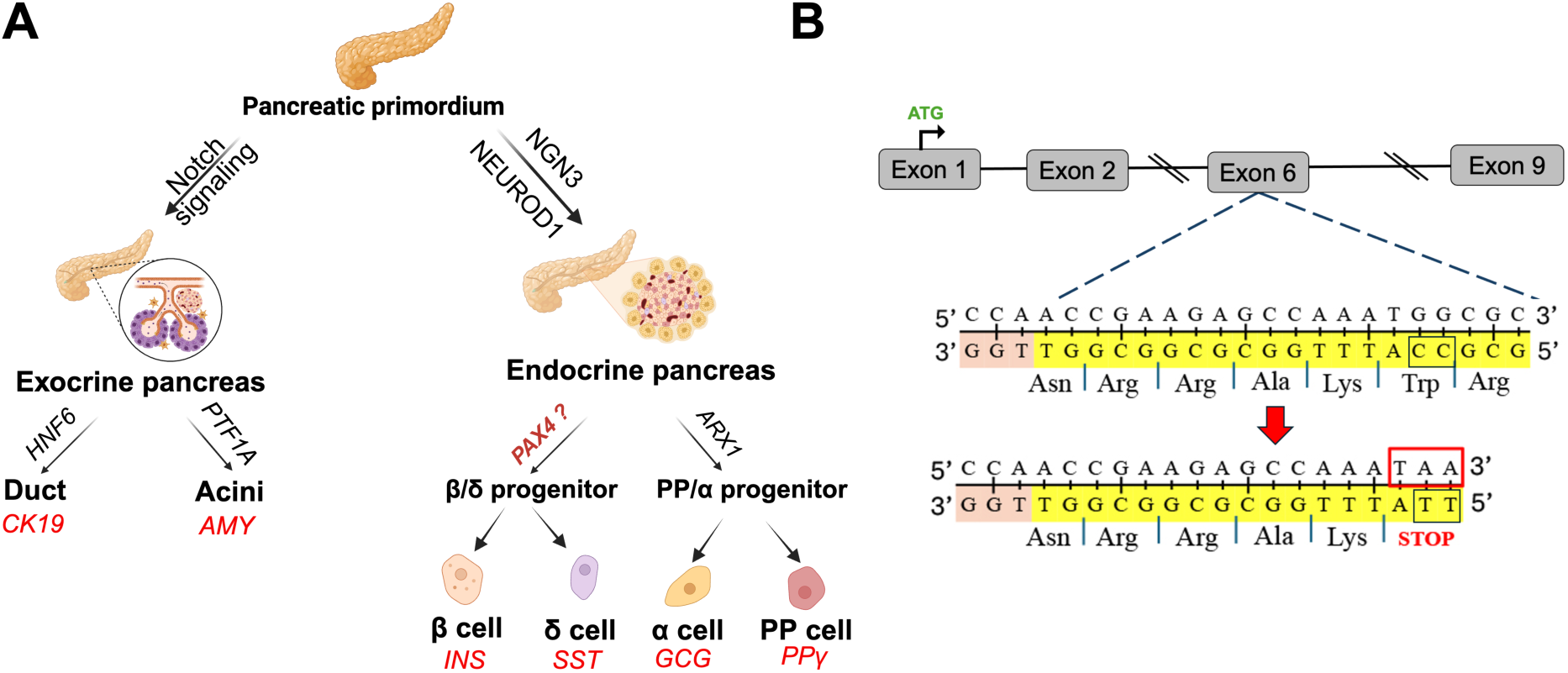
Schematic overview of pancreatic lineage allocation and CRISPR design for functional inactivation of *PAX4*. A) Schematic of pancreatic progenitor differentiation highlighting *PAX4*-driven endocrine commitment. B) CRISPR/Cas9 target site and editing strategy for generating a PAX4 loss-of-function allele

Although rodent models have been essential for elucidating core developmental pathways, significant interspecies differences limit their universal translational relevance. For example, human islets display distinct cytoarchitecture, cellular composition, and functional responses compared to murine islets (9–13). Furthermore, emerging evidence indicates that the timing and regulation of endocrine differentiation programs, including those involving *NGN3* and *PAX* genes, differ markedly between rodents and humans. These differences underscore the need for alternative model systems that bridge the gap between rodent genetics and human pancreatic biology (14–16). The pig serves as a valuable “bridge” model for investigating pancreatic development and diabetes. Porcine pancreas development exhibits anatomical, physiological, and molecular similarities to that of humans, including comparable islet size, vascularization, and endocrine cell organization (17, 18). Pig islets have been considered as a promising source for xenotransplantation, and recent advances in genome editing enable precise genetic manipulation of key developmental regulators in this species (19). Despite the recognized translational potential of pigs, the role of core endocrine fate determinants, such as *PAX4*, has not been systematically examined in porcine pancreatic development.

Recent advances in CRISPR/Cas9-based genome editing, including base-editing technologies, have enabled efficient and precise generation of loss-of-function alleles in large animals without introducing double-strand breaks (20). These methods offer a robust platform for modeling human disease-associated variants and investigating gene function in physiologically relevant contexts. Utilizing these tools, this study will investigate whether porcine models can be engineered to investigate whether developmental paradigms identified in rodents are conserved in large mammals; and whether the pigs can serve as a robust translational model for investigating human-relevant genetic mechanisms of diabetes and provide new insights into the role of *PAX4* in pancreatic endocrine architecture and function.

## Materials and Methods

### Ethics Statement

All experiments involving live animals were conducted in accordance with the guidelines approved by the University of Missouri Institutional Animal Care and Use Committee (IACUC protocol #45081).

### Generation of PAX4 knockout piglets

Guide RNA design and base editor preparation: Three candidate single-guide RNAs (sgRNAs) targeting exons 2 and 6 of the porcine *PAX4* locus were designed using the BE-Designer online tool (http://www.rgenome.net) and synthesized by IDT (Fig. 1B; Fig. S1). BE4 mRNA was produced via *in vitro* transcription of the pCMV-BE4-RrA3F plasmid (Addgene #138340; a gift from Nicole Gaudelli) using the mMESSAGE mMACHINE T7 kit (Invitrogen) according to the manufacturer’s protocol.

### *In vitro* oocyte maturation, parthenogenetic activation, and embryo culture

Unless otherwise specified, all chemicals were purchased from Sigma-Aldrich (St. Louis, MO). *In vitro* maturation (IVM), *in vitro* fertilization (IVF), and embryo culture were performed according to established protocols (21–23). Cumulus-oocyte complexes (COCs) were collected from abattoir-derived sow ovaries (ART Inc., Madison, WI; or DeSoto Biosciences, Seymour, TN) and transported overnight in commercial maturation medium #1 at 38.5 °C. After 24 hours, groups of 50-75 COCs were transferred to maturation medium #2 and cultured for an additional 20 hours at 38.5 °C in 5% CO₂ and 100% humidity. Upon completion of maturation, cumulus cells were removed by vortexing COCs in 0.1% hyaluronidase prepared in HEPES-buffered medium containing 0.01% polyvinyl alcohol (PVA) for 4 minutes.

### Parthenogenetic embryos

For parthenogenetic activation, denuded oocytes were placed in an activation medium (280 mM mannitol, 0.1 mM CaCl₂, 0.05 mM MgCl₂, 0.5 mM HEPES, 0.1% BSA) between platinum electrodes and activated using two direct-current pulses (1 kV/cm, 30 µs) delivered by a Super electro cell fusion generator (Nepa Gene, Japan). Following a 2-min equilibration, embryos were incubated for 4 h in PZM3 medium supplemented with 2 mM 6-dimethylaminopurine.

### In vitro fertilization

Groups of 50 mature oocytes were placed in 500 µL modified Tris-buffered medium (mTBM) in four-well culture dishes. Fresh extended boar semen (1 mL) was diluted to 10 mL with Dulbecco’s phosphate-buffered saline (DPBS) containing 1 mg/mL BSA and centrifuged at 1,000 x g for 4 min at 25 °C. After three washes, spermatozoa were resuspended in mTBM and added to oocytes at a final concentration of 5 × 10⁵ sperm/mL. Gametes were co-incubated for 5 h at 38.5 °C and 5% CO2. Presumptive zygotes were subsequently cultured in PZM3 supplemented with 3 mg/mL fatty acid-free BSA at 38.5 °C in 5% CO2, 5% O2, and 100% humidity.

### Embryo microinjection and embryo transfer

Parthenogenetic or IVF-derived embryos were microinjected with a mixture of RrA3F-BE4 mRNA and target sgRNA using a FemtoJet microinjector (Eppendorf). BE4 mRNA (500 ng/µL) and sgRNA (2 µM) were combined and diluted in RNase-free water, then diluted to final concentrations of 50 ng/µL for BE4 mRNA and 0.2 µM for sgRNA for injection. Parthenogenetic embryos that were injected were cultured to the blastocyst stage (144 hours) to assess editing efficiency (Fig. S1). Injected IVF embryos were surgically transferred into a single oviduct of synchronized recipient sows to produce live offspring. For each embryo transfer, 70 embryos were injected with BE4 mRNA and *PAX4* sgRNA, and 30 control embryos were injected with the same injection buffer but without BE4/sgRNA, and both sets were co-transferred into the same recipient to establish *PAX4* knockout (KO) and control wild-type (WT) piglets.

### Genotyping and editing analysis

Individual blastocysts were washed in PBS-PVA and lysed in Direct PCR lysis buffer (Vivagen) containing proteinase K (100 µg/mL) at 65 °C for 1 h, followed by enzyme inactivation at 95°C for 10 min. Tissue biopsies (ear notch or tail dock) and fetal tissues were digested overnight at 55°C in lysis buffer containing Tris-HCl, NaCl, EDTA, SDS, RNase A, and proteinase K. Genomic DNA was extracted using phenol-chloroform, ethanol-precipitated, and resuspended in 10 mM Tris-HCl (pH 7.4). DNA quantity and purity were assessed using a NanoDrop 8000 spectrophotometer. Targeted editing at the PAX4 locus was evaluated using nested PCR with locus-specific primers (Table S1). PCR products were resolved on 2% agarose gels, purified using CleanNGS magnetic beads (CleanNA), and subjected to Sanger sequencing. Chromatograms were analyzed using SnapGene v7.2.0 to confirm on-target base editing.

### *In silico* protein structure modeling

Three-dimensional structures of WT and KO porcine PAX4 proteins were predicted using ColabFold, an AlphaFold2-based implementation (24). The WT reference PAX4 amino acid sequence (NCBI accession XP_020934396.1) served as input, while the KO sequence was generated by truncation downstream of the premature stop codon. Structure prediction used MMseqs2-based multiple sequence alignment, and top-ranked models were selected based on pLDDT confidence scores. Structural refinement was conducted using AMBER relaxation, and the resulting PDB files were used for comparative analysis.

### Animal procedures and sample collection

Piglets were weighed daily after birth, and body condition was assessed using a standardized 5-point scoring system. Whole blood was collected by venipuncture into serum tubes, allowed to clot, centrifuged at 2,000 x g for 15 minutes, and serum was stored at-80°C. Serum biochemistry analyses were conducted by the University of Missouri Veterinary Medical Diagnostic Laboratory. Pancreatic tissue was collected immediately following euthanasia, rinsed in cold PBS, and divided into two sections for downstream applications. One portion was snap-frozen for snRNA sequencing, and the remaining tissue was fixed in 4% paraformaldehyde overnight, transferred to 70% ethanol, and processed for paraffin embedding. Sections (5 µm) were prepared for histological and immunostaining analyses.

### Immunofluorescence and immunohistochemistry

Paraffin sections were deparaffinized, rehydrated, and subjected to heat-mediated antigen retrieval in citrate buffer (pH 6.0). Sections were permeabilized with 1% Triton X-100 and blocked using SuperBlock™ or 5% BSA. Primary antibodies against insulin, glucagon, and somatostatin were applied overnight at 4°C, followed by fluorophore-conjugated secondary antibodies (Table S2). Nuclei were counterstained with DAPI, and sections were mounted using antifade medium. For insulin immunohistochemistry, endogenous peroxidase activity was quenched prior to antibody incubation. Detection was performed using HRP-conjugated secondary antibodies and DAB substrate, followed by hematoxylin counterstaining. Images were captured using a Leica DM4000B microscope with consistent exposure settings across genotypes.

### SnRNA Sequencing

#### Library preparation and sequencing

Nuclei were isolated from frozen pancreatic tissue using the 10x Genomics Chromium Nuclei Isolation Kit. Single-nucleus gene expression libraries were prepared following the 10x Chromium workflow and sequenced as paired-end reads. In total, four biological replicates per genotype were processed (4 WT and 4 KO).

#### Reference construction and read processing

A pre-mRNA reference genome was generated from Ensembl release 111 (Sscrofa11.1) to include intronic reads. FASTQ quality was assessed using fastp, and alignment metrics were evaluated using samtools (25). Reads were processed using Cell Ranger v8.0.1 to generate filtered gene-barcode matrices.

#### Seurat analysis and Cell-type annotation

Filtered matrices were imported into Seurat (R). Nuclei were filtered, normalized, and clustered using standard workflows. Dimensionality reduction was performed using PCA and UMAP. Cell types were annotated based on canonical pancreatic markers, and genotype-specific differences were assessed by comparing cluster composition and marker expression.

### Bulk RNA sequencing

Bulk RNA-seq was performed using RNA isolated from frozen neonatal pancreatic tissue. In total, three biological replicates per genotype were analyzed (3 WT and 3 KO). Libraries were prepared using the Plasmidsaurus stranded 3′ end-counting RNA-seq workflow, in which polyadenylated mRNA is reverse-transcribed using oligo-dT priming, followed by second-strand synthesis, tagmentation, indexing, and amplification. Unique molecular identifiers were incorporated to remove PCR duplicates, and libraries were sequenced on an Illumina platform as single-end reads.

Sequencing data were processed to generate quality-control outputs, aligned deduplicated BAM files, and gene-level count matrices. Because this workflow quantifies reads from the 3′ end of transcripts, the data were interpreted at the gene-expression level rather than for transcript isoform analysis. Gene-count matrices were imported into R, and differential expression analysis between KO and WT pancreas was performed using DESeq2. Selected endocrine, pancreatic, and diabetes-associated genes were visualized by targeted heatmap, and GO enrichment analysis of genes upregulated in KO pancreas was performed using clusterProfiler.

## Statistical analysis

All statistical analyses were performed in R (v4.5.0) (Table S3). Parametric and nonparametric tests were selected based on data distribution. Survival was analyzed using Kaplan-Meier curves and log-rank tests. Serum chemistry values are reported as median (Q1, Q3). A two-sided p < 0.05 was considered statistically significant.

## Data availability

A total of 8 snRNA-seq data sets generated for this article can be accessed via NCBI Sequence Read Archive (SRA) Bioprojects with the accession number PRJNA1415818.

## Results

### Functional inactivation of *PAX4* resulted in viable but metabolically compromised piglets

Three candidate sgRNAs were designed to introduce an in-frame premature translational stop codon within the *PAX4* coding sequence. One sgRNA exhibited both high embryo developmental competence and robust on-target editing efficiency and was selected for all subsequent experiments (Fig. S1). Introduction of the stop codon in the *PAX4* coding sequence at the target site results in the loss of crucial DNA-binding homeobox domain, rendering the resulting protein nonfunctional (Fig. S2). Following *in vitro* validation, IVF embryos were microinjected with BE4/sgRNA alongside sham-injected controls, and transferred into two estrus synchronized recipients, resulting in one successful pregnancy and birth of 14 live piglets. Genotyping confirmed five WT and nine *PAX4* KO piglets (Fig. 2A; Fig. S3).

**Figure 2.**
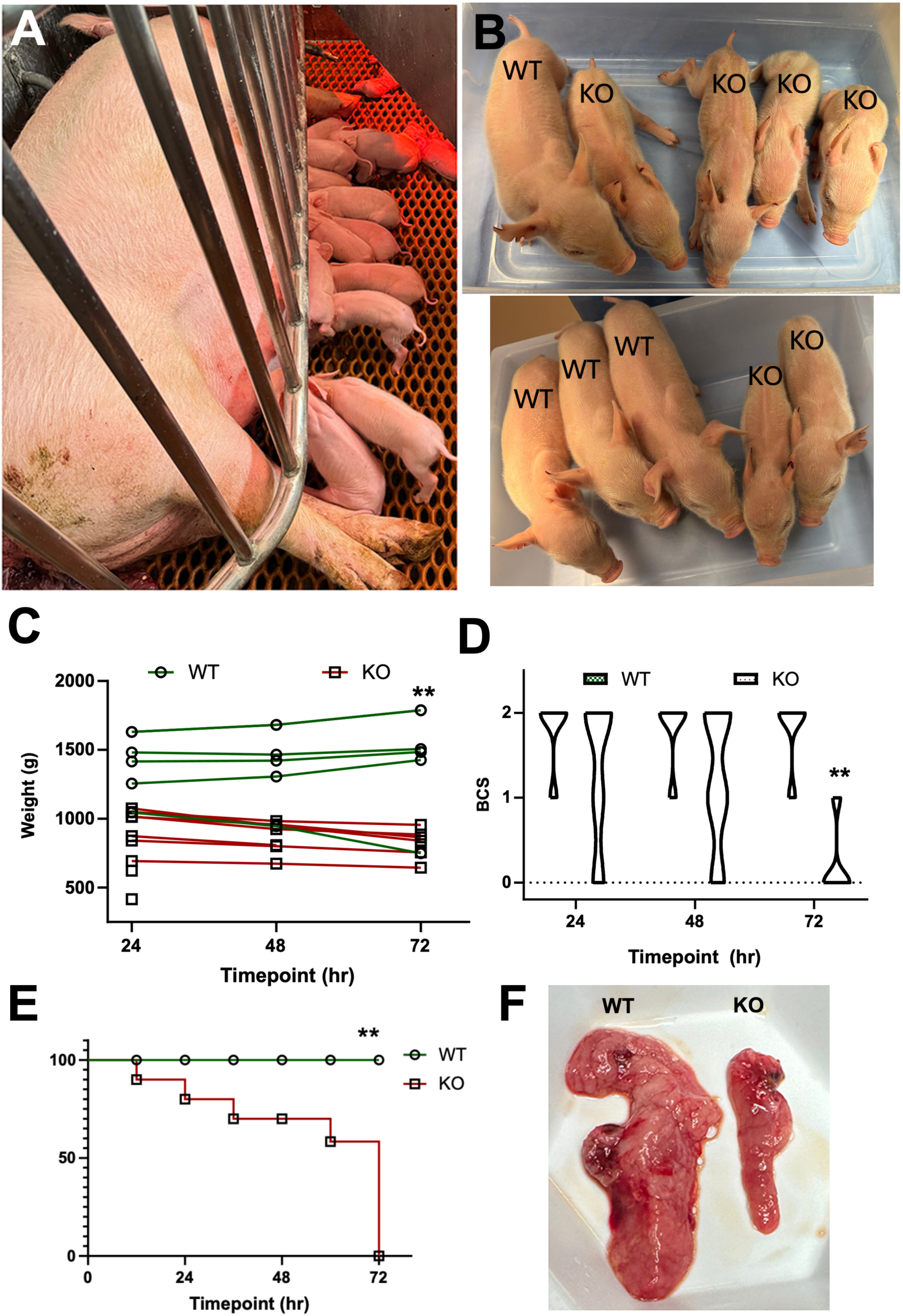
Longitudinal assessment of body weight, body condition, glycemia, and survival in WT and KO piglets. A) Litter of WT and *PAX4*-KO piglets with the sow. B) Gross comparison of the size of WT vs KO piglets. C) Body weight, D) Body condition score (BCS), and E) Kaplan-Meier survival curve of WT and KO piglets. F) Comparative morphology of pancreas from WT and KO piglets at necropsy.

At birth, KO piglets were visibly smaller than WT littermates and exhibited a rapid decline in growth and clinical status during the early postnatal period (Fig. 2B). Quantitative analysis demonstrated a progressive and significant reduction in body weight among KO piglets during the first 72 hours postnatally (p < 0.001), while WT piglets maintained or gained weight over the same period (Fig. 2C). Correspondingly, body condition scores (BCS) declined significantly in KO animals (p < 0.001), with several reaching critically low values by 72 hours (Fig. 2D). This deterioration was accompanied by neurological manifestations such as ataxia, incoordination, and seizures. Survival analysis revealed a significant genotype-dependent difference in early postnatal viability (log-rank test, p = 0.033). All WT piglets survived the 72-hour observation period, whereas KO piglets experienced progressive mortality beginning within the first 24 hours, with none surviving beyond 72 hours (Fig. 2E). Gross pathological examination at necropsy showed a visibly reduced pancreatic mass in KO piglets compared to WT littermates (Fig. 2F). The WT pancreata were well developed with broad, lobulated morphology, while KO pancreata were consistently smaller and less lobulated, indicating impaired pancreatic development.

### *PAX4* deficiency causes rapid metabolic decompensation and multisystem biochemical abnormalities

Serum biochemical profiling indicated rapid and severe metabolic deterioration in *PAX4*-KO piglets during the first 72 hours of life (Table 1; Fig. 3). In line with point-of-care glucose measurements, KO piglets developed pronounced hyperglycemia within the first day, which progressively worsened over time (Fig. 3A). By postnatal day 3, indicators of renal dysfunction were evident, with significantly elevated blood urea nitrogen (BUN) and creatinine levels in KO piglets compared to WT controls. Lipid metabolism was also severely disrupted, as shown by increased circulating triglycerides (Fig. 3B), total cholesterol (Fig. 3C), and non-esterified fatty acids (NEFA; Fig. 3D). Consistent with renal dysfunction, electrolyte homeostasis was markedly disrupted in the KO piglets, with significant hypernatremia and hyperchloremia, increased serum osmolality (Fig. 3E), and an elevated anion gap (Fig. 3F), resulting in metabolic acidosis. In contrast, total and direct bilirubin levels did not differ significantly between genotypes (p > 0.05), indicating preserved hepatic bilirubin handling at this stage.

**Figure 3.**
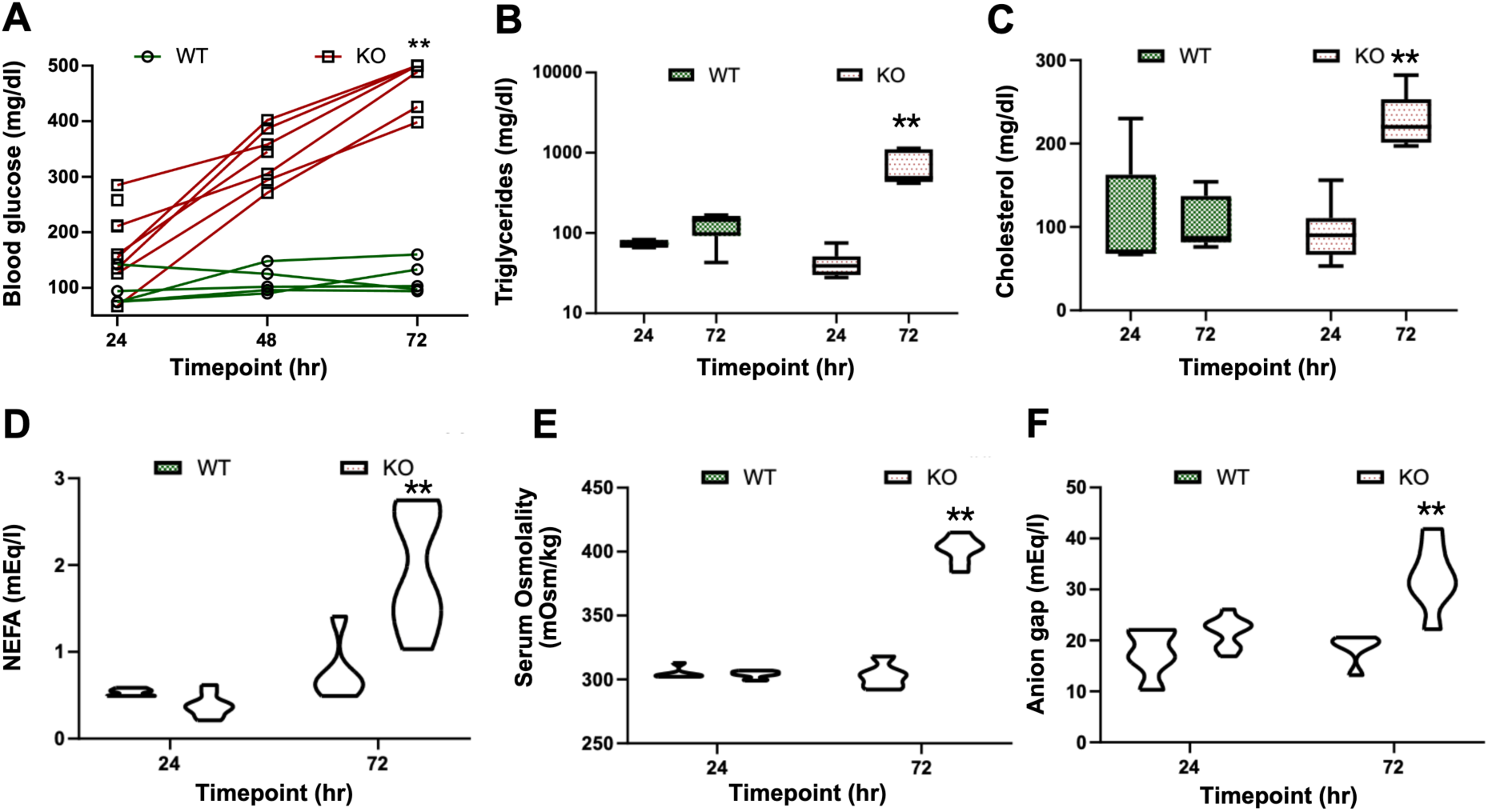
Metabolic profiling of WT and PAX4-KO neonatal piglets at 24 and 72 h post-birth. (A) Blood glucose; (B) triglycerides; (C) cholesterol; (D) NEFA; (E) serum osmolality; and (F) anion gap measurements in WT and KO piglets. Blood glucose and triglyceride concentrations are plotted on a log₁₀ scale to accommodate their wide dynamic range. Each plot displays data as a combination of box and violin elements, where boxes indicate the interquartile range (IQR) with horizontal lines marking the median, and violins represent the distribution density.

**Table 1.** Serum biochemical profile of WT and KO piglets on Day 1 and Day 3.

| S. No. | Component | Day 1 |  | Day 3 |  | P-values | Chi square |
| --- | --- | --- | --- | --- | --- | --- | --- |
|  |  | WT | KO | WT | KO |  |  |
| 1 | Glucose (mg/dL) | 61<sup>a</sup><br>(61, 71) | 172<sup>b</sup><br>(146, 372) | 96<sup>a</sup><br>(89, 101) | **728<sup>b</sup>**<br>(686, 902) | 0.001 | 22.855 |
| 2 | BUN(mg/dL) | 16<sup>a</sup><br>(15, 17) | 18<sup>a</sup><br>(17, 20) | 12<sup>a</sup><br>(11, 13) | **71<sup>b</sup>**<br>(65.5, 77) | 0.001 | 18.732 |
| 3 | Creatinine (mg/dL) | 1.2<sup>b</sup><br>(1.2, 1.4) | 1.1<sup>b</sup><br>(1.1, 1.3) | 0.4<sup>a</sup><br>(0.4, 0.6) | 1.2<sup>b</sup><br>(1.0, 1.35) | 0.001 | 12.788 |
| 4 | Sodium (mEq/L) | 128<sup>a</sup><br>(118,128) | 125<sup>a</sup><br>(124, 127) | 133<sup>ab</sup><br>(132,134) | 151<sup>b</sup><br>(148, 154) | 0.001 | 20.533 |
| 5 | Potassium (mEq/L) | 5.6<sup>b</sup><br>(5.5,5.8) | 5.6<sup>b</sup><br>(5.1,5.8) | 4.2<sup>b</sup><br>(3.9,4.6) | 5.5<sup>b</sup><br>(5.2,5.8) | 0.01 | 11.677 |
| 6 | Chloride (mEq/L) | 96<sup>a</sup><br>(95, 97) | 92<sup>a</sup><br>(91, 93) | 96<sup>a</sup><br>(95, 96) | 104<sup>b</sup><br>(102, 106) | 0.001 | 19.137 |
| 7 | Albumin (g/dL) | 1.2<sup>a</sup><br>(1.2, 1.4) | 1.1<sup>a</sup><br>(1.0, 1.2) | 1.4<sup>b</sup><br>(1.3, 1.5) | 1.2<sup>a</sup><br>(1.2, 1.35) | 0.05 | 8.581 |
| 8 | Total protein (g/dL) | 3.9<sup>b</sup><br>(3.4, 4.3) | 3.1<sup>a</sup><br>(2.6, 3.5) | 5.5<sup>b</sup><br>(4.9, 5.7) | 5.6<sup>b</sup><br>(5.25,6.15) | 0.001 | 17.394 |
| 9 | Globulin (g/dL) | 2.5<sup>ab</sup><br>(2.2, 2.7) | 1.9<sup>a</sup><br>(1.6, 2.5) | 4.0<sup>b</sup><br>(3.6, 4.5) | 4.3<sup>b</sup><br>(4.0, 4.9) | 0.001 | 19.203 |
| 10 | Triglyceride (mg/dL) | 39<sup>a</sup><br>(30, 40) | 69<sup>a</sup><br>(69, 81) | 145<sup>a</sup><br>(142, 161) | **487<sup>b</sup>**<br>(432,1105) | 0.001 | 21.010 |
| 11 | Cholesterol (mg/dL) | 70<sup>a</sup><br>(70, 95) | 90<sup>a</sup><br>(70, 108) | 87<sup>a</sup><br>(87, 120) | **220<sup>b</sup>**<br>(203, 238) | 0.010 | 12.811 |
| 12 | Total Bilirubin (mg/dL) | 1.1<br>(1.1, 1.2) | 1.0<br>(0.9,1.1) | 0.8<br>(0.7, 0.8) | 0.6<br>(0.5, 0.95) | >0.05 | 8.409 |
| 13 | Direct Bilirubin (mg/dL) | 0.0<br>(0.0, 0.0) | 0.0<br>(0.0, 0.0) | 0.1<br>(0.1, 0.1) | 0<br>(0.0, 0.1) | >0.05 | 6.847 |
| 14 | NEFA (mEq/L) | 0.54<sup>a</sup><br>(0.49,0.56) | 0.36<sup>a</sup><br>(0.31, 0.42) | 0.68<sup>a</sup><br>(0.60,0.69) | **1.74<sup>b</sup>**<br>(1.44,2.52) | 0.001 | 19.496 |
| 15 | Osmolality (mOsm/Kg) | 303<sup>a</sup><br>(303, 303) | 304<sup>a</sup><br>(303, 306) | 304<sup>a</sup><br>(296, 304) | **403<sup>b</sup>**<br>(396, 407) | 0.001 | 15.213 |
| 16 | Anion gap (mEq/L) | 16.8<sup>a</sup><br>(16.5,21.6) | 22.5<sup>a</sup><br>(18.8, 23.2) | 19.8<sup>a</sup><br>(18.9,19.8) | **32.1<sup>b</sup>**<br>(29.5,35.8) | 0.001 | 15.664 |

WT: Wild-type; KO: PAX4 knockout; BUN: blood urea nitrogen; NEFA: non-esterified fatty acids.
Data are presented as median (Q1, Q3)

**Table 1. Serum biochemical profile of WT and KO piglets on Day 1 and Day 3**
| S. No. | Component | Day 1 |  | Day 3 |  | P-values | Chi square |
| --- | --- | --- | --- | --- | --- | --- | --- |
|  |  | WT | KO | WT | KO |  |  |
| 1 | Glucose (mg/dL) | 61<sup>a</sup><br>(61, 71) | 172<sup>b</sup><br>(146, 372) | 96<sup>a</sup><br>(89, 101) | **728<sup>b</sup>**<br>(686, 902) | 0.001 | 22.855 |
| 2 | BUN(mg/dL) | 16<sup>a</sup><br>(15, 17) | 18<sup>a</sup><br>(17, 20) | 12<sup>a</sup><br>(11, 13) | **71<sup>b</sup>**<br>(65.5, 77) | 0.001 | 18.732 |
| 3 | Creatinine (mg/dL) | 1.2<sup>b</sup><br>(1.2, 1.4) | 1.1<sup>b</sup><br>(1.1, 1.3) | 0.4<sup>a</sup><br>(0.4, 0.6) | 1.2<sup>b</sup><br>(1.0, 1.35) | 0.001 | 12.788 |
| 4 | Sodium (mEq/L) | 128<sup>a</sup><br>(118, 128) | 125<sup>a</sup><br>(124, 127) | 133<sup>ab</sup><br>(132, 134) | 151<sup>b</sup><br>(148, 154) | 0.001 | 20.533 |
| 5 | Potassium (mEq/L) | 5.6<sup>b</sup><br>(5.5, 5.8) | 5.6<sup>b</sup><br>(5.1, 5.8) | 4.2<sup>b</sup><br>(3.9, 4.6) | 5.5<sup>b</sup><br>(5.2, 5.8) | 0.01 | 11.677 |
| 6 | Chloride (mEq/L) | 96<sup>a</sup><br>(95, 97) | 92<sup>a</sup><br>(91, 93) | 96<sup>a</sup><br>(95, 96) | 104<sup>b</sup><br>(102, 106) | 0.001 | 19.137 |
| 7 | Albumin (g/dL) | 1.2<sup>a</sup><br>(1.2, 1.4) | 1.1<sup>a</sup><br>(1.0, 1.2) | 1.4<sup>b</sup><br>(1.3, 1.5) | 1.2<sup>a</sup><br>(1.2, 1.35) | 0.05 | 8.581 |
| 8 | Total protein (g/dL) | 3.9<sup>b</sup><br>(3.4, 4.3) | 3.1<sup>a</sup><br>(2.6, 3.5) | 5.5<sup>b</sup><br>(4.9, 5.7) | 5.6<sup>b</sup><br>(5.25, 6.15) | 0.001 | 17.394 |
| 9 | Globulin (g/dL) | 2.5<sup>ab</sup><br>(2.2, 2.7) | 1.9<sup>a</sup><br>(1.6, 2.5) | 4.0<sup>b</sup><br>(3.6, 4.5) | 4.3<sup>b</sup><br>(4.0, 4.9) | 0.001 | 19.203 |
| 10 | Triglyceride (mg/dL) | 39<sup>a</sup><br>(30, 40) | 69<sup>a</sup><br>(69, 81) | 145<sup>a</sup><br>(142, 161) | **487<sup>b</sup>**<br>(432, 1105) | 0.001 | 21.010 |
| 11 | Cholesterol (mg/dL) | 70<sup>a</sup><br>(70, 95) | 90<sup>a</sup><br>(70, 108) | 87<sup>a</sup><br>(87, 120) | **220<sup>b</sup>**<br>(203, 238) | 0.010 | 12.811 |
| 12 | Total Bilirubin (mg/dL) | 1.1<br>(1.1, 1.2) | 1.0<br>(0.9, 1.1) | 0.8<br>(0.7, 0.8) | 0.6<br>(0.5, 0.95) | >0.05 | 8.409 |
| 13 | Direct Bilirubin (mg/dL) | 0.0<br>(0.0, 0.0) | 0.0<br>(0.0, 0.0) | 0.1<br>(0.1, 0.1) | 0<br>(0.0, 0.1) | >0.05 | 6.847 |
| 14 | NEFA (mEq/L) | 0.54<sup>a</sup><br>(0.49, 0.56) | 0.36<sup>a</sup><br>(0.31, 0.42) | 0.68<sup>a</sup><br>(0.60, 0.69) | **1.74<sup>b</sup>**<br>(1.44, 2.52) | 0.001 | 19.496 |
| 15 | Osmolality (mOsm/Kg) | 303<sup>a</sup><br>(303, 303) | 304<sup>a</sup><br>(303, 306) | 304<sup>a</sup><br>(296, 304) | **403<sup>b</sup>**<br>(396, 407) | 0.001 | 15.213 |
| 16 | Anion gap (mEq/L) | 16.8<sup>a</sup><br>(16.5, 21.6) | 22.5<sup>a</sup><br>(18.8, 23.2) | 19.8<sup>a</sup><br>(18.9, 19.8) | **32.1<sup>b</sup>**<br>(29.5, 35.8) | 0.001 | 15.664 |

### SnRNA sequencing reveals selective loss of endocrine lineages following *PAX4* inactivation

SnRNA-seq was conducted on pancreatic tissue from WT and KO piglets to characterize the cellular consequences of *PAX4* loss. Initial Cell Ranger outputs identified approximately 90,000 barcodes per sample prior to quality control. After Seurat filtering, substantially fewer high-quality nuclei were retained in the KO dataset, indicating reduced cellular complexity. Total UMI counts and the number of detected genes were strongly correlated (r = 0.88), supporting the high quality of the snRNA-seq data. WT nuclei exhibited greater transcript complexity, while KO nuclei were predominantly concentrated in low-count, low-feature regions (Fig. S4A, C). P-based clustering resolved the major pancreatic compartments, including acinar, endocrine, stromal, endothelial, and immune populations (Fig. 4A). Acinar cells constituted the majority of nuclei in both genotypes, while stromal and endothelial cells formed smaller, distinct clusters. Within the endocrine compartment, however, a striking genotype-dependent shift was observed. WT pancreata contained well-defined α-, β-, and δ-cell populations, whereas KO endocrine nuclei were markedly reduced and consisted almost exclusively of α-cells (Fig. 4B).

**Figure 4.**
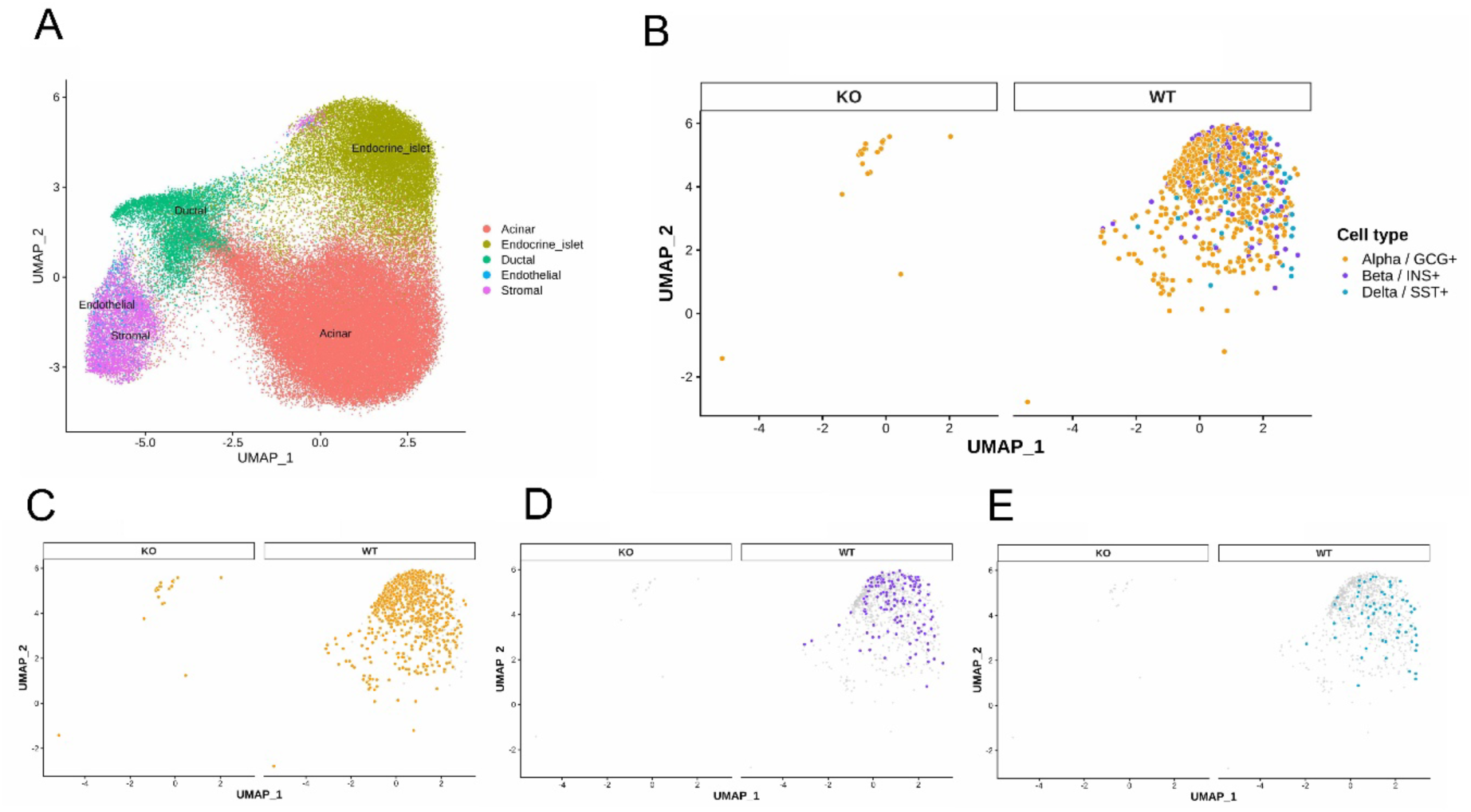
Single-nucleus RNA-seq UMAPs of neonatal pig pancreas identify major cell populations and reveal endocrine cell loss in *PAX4*-KO. (A) Integrated UMAP showing major pancreatic cell types. (B) UMAP of endocrine nuclei with α-, β-, and δ-cell identities shown separately for WT and *PAX4*-KO (C–E) Genotype-split UMAPs highlighting marked depletion of α-, δ-, and β-cells in *PAX4*-KO compared with WT.

Feature-level expression maps confirmed the preservation of glucagon-expressing α-cells in KO pancreata (Fig. 4C), whereas insulin-expressing β-cells (Fig. 4D) and somatostatin-expressing δ-cells (Fig. 4E) were undetectable. Dot-plot visualization of canonical marker genes further validated these cell-type assignments (Fig. S4B).

### Bulk RNA-seq validates endocrine marker loss and reveals compensatory transcriptional remodelling in *PAX4*-KO pancreas

Bulk RNA-seq analysis further supported the snRNA-seq evidence of endocrine-lineage disruption in *PAX4*-KO neonatal pancreas. Differential expression analysis showed marked suppression of key endocrine markers, particularly *INS,* consistent with loss of β-cell identity, while several genes associated with metabolic stress and diabetes-related transcriptional responses, including *SORD*, *TXNIP*, *IGF2*, and *MT1A*, were increased in KO pancreas (Fig. S5A). A targeted heatmap of endocrine, pancreatic, and diabetes-associated genes confirmed the reduction of β-and δ-cell markers such as *INS* and *SST*, while also highlighting genotype-associated shifts in selected endocrine and stress-responsive genes across WT and KO samples (Fig. S5B).

To characterize the biological programs associated with genes upregulated in *PAX4*-KO pancreas, GO enrichment analysis was performed. Upregulated genes were enriched for biological processes related to epithelial differentiation, tissue development, digestive system development, protein sumoylation, histone methylation, and xenobiotic response (Fig. S6A). Molecular function and cellular component enrichment further indicated increased representation of actin/cytoskeletal binding terms and structural components such as sarcomere, myofibril, contractile fiber, adherens junction, and actin cytoskeleton (Fig. S6B, C), suggesting that *PAX4* loss is accompanied not only by endocrine cell depletion but also by broader remodeling of pancreatic developmental, epithelial, and cytoskeletal transcriptional programs.

### Immunofluorescence confirms selective loss of β- and δ-cells with preservation of exocrine architecture

Immunofluorescence analyses supported the snRNA-seq findings and provided histological confirmation of altered endocrine composition. Amylase staining showed comparable distribution and intensity in WT and KO pancreata, with well-organized acinar structures in both genotypes, indicating preservation of exocrine architecture following *PAX4* inactivation (Fig. 5). In contrast, endocrine staining patterns differed markedly between genotypes. WT pancreatic sections displayed prominent islet clusters containing insulin-, glucagon-, and somatostatin-positive cells, with merged images revealing intact islet architecture comprising α-, β-, and δ-cell populations (Fig. 6; Figs. S7, S8 and S9). In KO pancreata, insulin and somatostatin signals were greatly reduced or absent, while glucagon remained the dominant endocrine hormone signal (Fig. 6; Figs. S7, S8 and S9). In summary, these data demonstrate that functional inactivation of *PAX4* in pigs leads to selective loss of β- and δ-cell lineages, increase of α-cells, intact exocrine development, and rapid postnatal metabolic failure.

**Figure 5.**
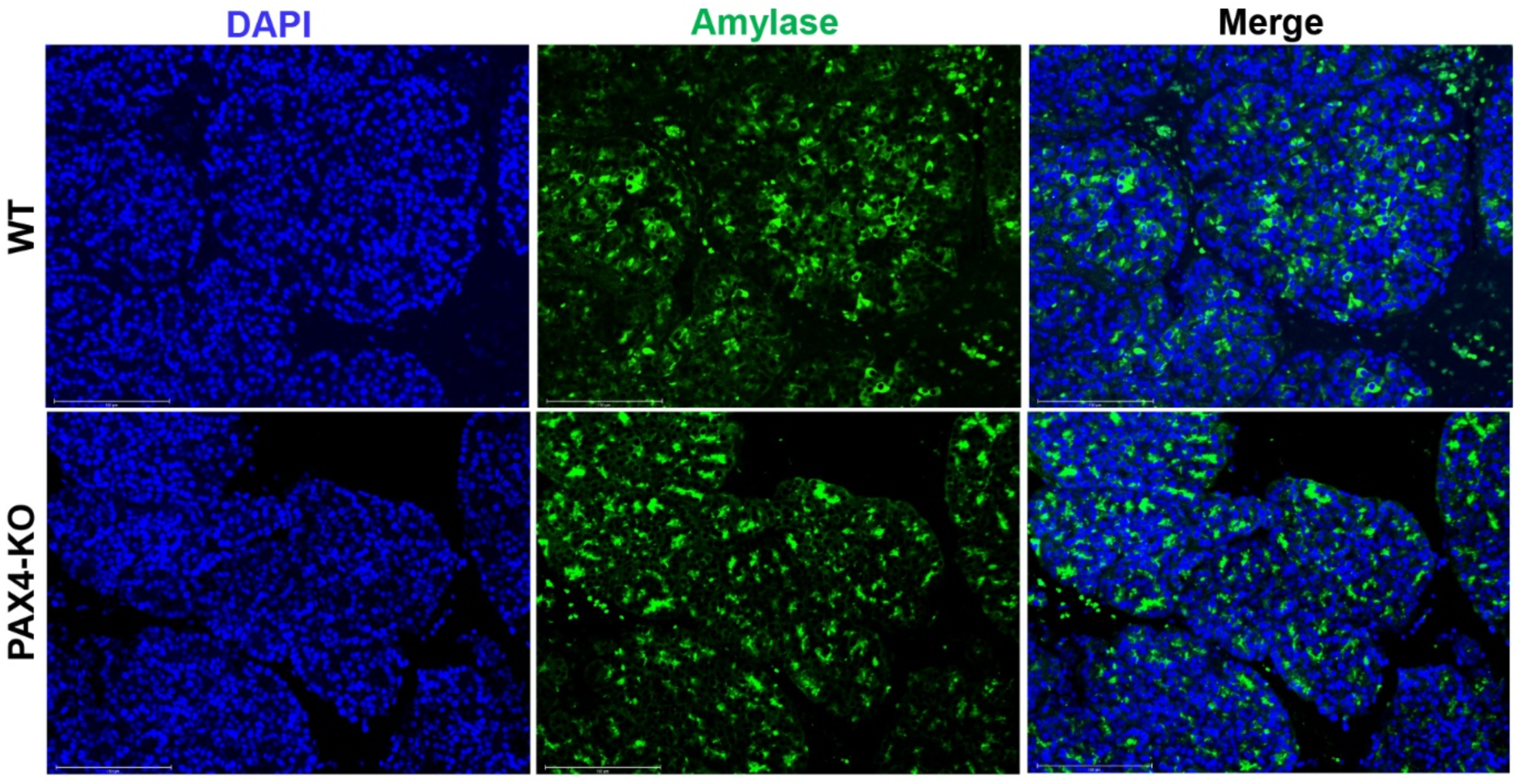
Immunofluorescence of acinar marker amylase in WT and KO pancreas showing preserved exocrine compartment in *PAX4*-KO piglets. Measuring bar: 150 mM

**Figure 6.**
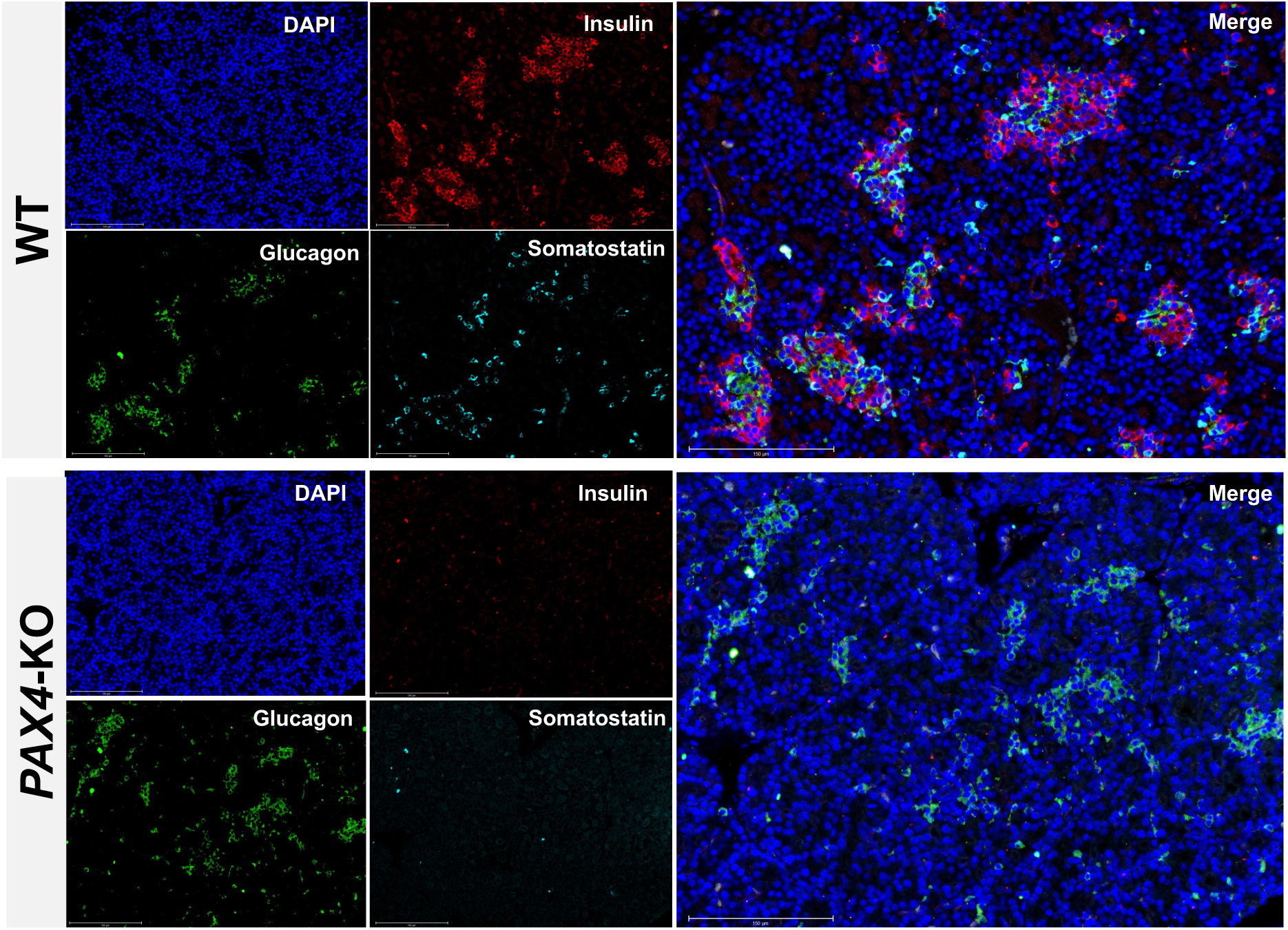
Multiplex immunofluorescence of pancreatic islets in WT and KO piglets showing loss of insulin and somatostatin with preserved glucagon signals. Measuring bar: 150 mM

## Discussion

In this study, we generated a *PAX4* loss-of-function (KO) pig model using CRISPR/Cas9 cytidine-deaminase base editing to introduce a targeted C-to-T substitution that creates an in-frame premature stop codon, resulting in a truncated, nonfunctional PAX4 protein. To our knowledge, this represents the first *PAX4*-deficient model in swine. *PAX4* is a critical gatekeeper transcription factor that directs endocrine progenitors toward insulin-producing β-cells and somatostatin-producing δ-cells. The essential role of *PAX4* in endocrine lineage specification has been established primarily through foundational KO studies in mice, while variants in the human *PAX4* locus have been associated with both monogenic and complex forms of diabetes (8, 26). The porcine *PAX4* loss-of-function model described here provides a human-relevant *in vivo* platform for interrogating endocrine lineage allocation, postnatal metabolic adaptation, and systemic diabetic decompensation at the organismal scale. Unlike rodents, pigs share key features with humans, including islet cytoarchitecture, β-cell maturation dynamics, and neonatal glucose homeostasis, enabling direct, translationally meaningful interpretation of cell-type-specific vulnerabilities (17). When combined with our ability to generate genetically modified pig models at high efficiency, these models are now positioned to generate translationally relevant models. As shown in this study, the availability of base editors has been a game-changer in generating *PAX4*-KO pig models at high efficiency. From a round of embryo transfer, we established 9 KO and 5 WT piglets from BE4-injected and sham-injected control embryos, respectively.

The *PAX4*-KO piglets exhibited a rapid postnatal decline, with marked loss of body weight and body condition by postnatal day 3, accompanied by progressive neurologic signs necessitating humane euthanasia. This phenotype reflects a profound failure of postnatal energy homeostasis. In the absence of functional β-cell-derived insulin, neonatal tissues cannot effectively utilize circulating glucose, resulting in a severe catabolic state (27). Consistent with this, hyperglycemia was detected within 24 h of birth and worsened over time, while WT littermates remained normoglycemic. The genotype x time interaction confirms a rapidly progressive diabetic phenotype rather than secondary effects of dehydration or terminal decline. This trajectory parallels observations in *PAX4*-deficient mice and rabbits and underscores a conserved requirement for *PAX4* in postnatal glucose regulation. Notably, the first day of life represents a major metabolic transition from continuous placental nutrient supply to intermittent milk intake, requiring rapid insulin-mediated control of feeding or fasting cycles (28). The early onset hyperglycemia observed in KO piglets during this window supports a primary endocrine failure rather than a secondary consequence of dehydration or terminal decline. Survival analysis further demonstrated uniform early mortality in KO piglets, closely matching the clinical progression and supporting endocrine failure as the primary cause of lethality.

Gross examination of the pancreata revealed a markedly hypoplastic pancreas in *PAX4*-KO piglets. This reduction in pancreatic mass indicates a critical role for *PAX4* not only in endocrine differentiation, but also in establishing or maintaining pancreatic growth during late gestation and early neonatal life. Although exocrine tissue was preserved, the dramatic loss of endocrine cells likely disrupts local trophic and paracrine signalling required for normal pancreatic development. This phenotype aligns with the temporal expression pattern of *PAX4* during endocrine lineage differentiation and with prior reports demonstrating its role in endocrine cell survival and stress resistance. Biochemical profiling revealed changes consistent with severe neonatal diabetes and metabolic decompensation. *PAX4*-KO piglets developed extreme hyperglycemia, leading to elevated serum osmolality by day 3, consistent with glucose-driven osmotic diuresis, dehydration, and electrolyte loss. Associated hypernatremia, hyperchloremia, and relative hyperkalemia reflect dehydration, acidosis, and insulin deficiency-mediated electrolyte shifts (29). Neurologic signs observed clinically are consistent with hyperosmolar stress on the central nervous system (30; 31). Lipid profiling demonstrated progressive increases in triglycerides, cholesterol, and non-esterified fatty acids, reflecting loss of insulin-mediated suppression of hepatic VLDL production and adipose lipolysis (32). The widened anion gap is consistent with ketoacidosis driven by excessive hepatic ketogenesis (33). Collectively, these findings define a biochemical signature of fulminant neonatal diabetes.

SnRNA sequencing confirmed loss of β- and δ-cell populations with relative preservation of α-cells. Integrated clustering identified major pancreatic compartments, with KO samples enriched for stromal and endothelial nuclei among retained cells. Immunofluorescence aligned closely with transcriptomic patterns. The preserved amylase expression, indicating intact exocrine differentiation, but revealed a near-complete loss of insulin-positive β-cells and somatostatin-positive δ-cells in the KO pancreata. This mirrors findings from *Pax4*-deficient rodents, rabbits, and recent human genetic studies linking biallelic *PAX4* loss-of-function variants to neonatal diabetes (8, 34, 35). In contrast, glucagon-expressing α-cells were increased, consistent with the established role of *PAX4* in promoting β- and δ-cell fate while repressing α-cell lineage commitment (36). These findings support a model in which α-cell differentiation represents a default endocrine pathway in the absence of *PAX4*, and that *PAX4* is a stringent requirement for β- or δ-lineage allocation (37).

Bulk RNA-seq further supported the cellular findings obtained by snRNA-seq and immunofluorescence. The marked reduction of endocrine markers, particularly INS and SST, is consistent with the near-complete loss of β- and δ-cell populations in *PAX4*-KO pancreas, while preservation or relative enrichment of selected non-β-cell and stress-associated transcripts likely reflects the altered cellular composition of the KO pancreas rather than a purely cell-intrinsic transcriptional effect (38). The targeted heatmap therefore provides an orthogonal validation of endocrine-lineage depletion at the whole-tissue level. In parallel, GO enrichment of genes upregulated in KO pancreas indicated activation of pathways related to epithelial differentiation, tissue remodeling, cytoskeletal organization, and developmental programs, suggesting that PAX4 loss is accompanied by broader pancreatic remodeling secondary to failed endocrine specification (39, 40).

The selective loss of β- and δ-cells, preservation of α-cells, and rapid onset of neonatal diabetes observed in this model parallel clinical features reported in individuals carrying deleterious *PAX4* variants. Beyond disease modelling, this platform provides a potential translational framework to link genotype with phenotype, evaluate cell-replacement and gene-correction strategies, and benchmark emerging diabetes therapeutics in a large-animal model prior to clinical translation.

## Author Contributions

R.P. and B.P.T. conceived the project and designed the experiments; R.P. and K.P. performed CRISPR guide RNA validations, embryo injections, embryo transfers and generated *PAX4* KO pigs. R.P. and A.S. performed phenotyping of KO piglets; R.P. generated the libraries and performed transcriptomic analysis; R.P. wrote the initial draft; B.P.T. revised the manuscript based on the input from all authors. All authors approved the final draft for submission.

## Acknowledgements

This research was supported by funding from Congressionally Directed Medical Research Programs (CDMRP), Department of Defence, USA, Award # W81XWH-22-1-0017.

