## Supplementary file for "Loss of *PAX4* results in disrupted endocrine pancreas development and neonatal diabetes in pigs"

**This PDF file includes:**

Supplementary Figures (S1 to S7)

Figure S1. CRISPR/Cas9 gRNA design, embryo development outcomes, and sequence validation for targeting porcine *PAX4*.

Figure S2. Predicted 3D structures of wild-type and mutant porcine *PAX4* proteins.

Figure S3. Sanger sequencing of *PAX4* alleles in WT and BE4-edited piglets.

Figure S4. SnRNA-seq reveals endocrine-cell depletion and broad transcriptional disruption in *PAX4*-KO neonatal pancreas.

Figure S5. Bulk RNA-seq shows loss of endocrine-cell markers and activation of diabetes-associated transcriptional signatures in *PAX4* KO neonatal pancreas.

Figure S6. Gene ontology enrichment analysis of genes upregulated in PAX4-KO neonatal pancreas

Figure S7. Representative H&E histology and insulin immunohistochemistry of WT and *PAX4*-KO neonatal pancreas

Figure S8. Immunofluorescence staining of WT neonatal pancreas

Figure S9. Immunofluorescence of *PAX4*-KO pancreas

Supplementary Tables (S1-S3)

Table S1. Overview of Guide RNA Sequences and Screening Primers

Table S2. Antibody Panel and Working Dilutions

Table S3. Statistical analysis plan and outputs for phenotypic, survival, and serum biochemistry data

A

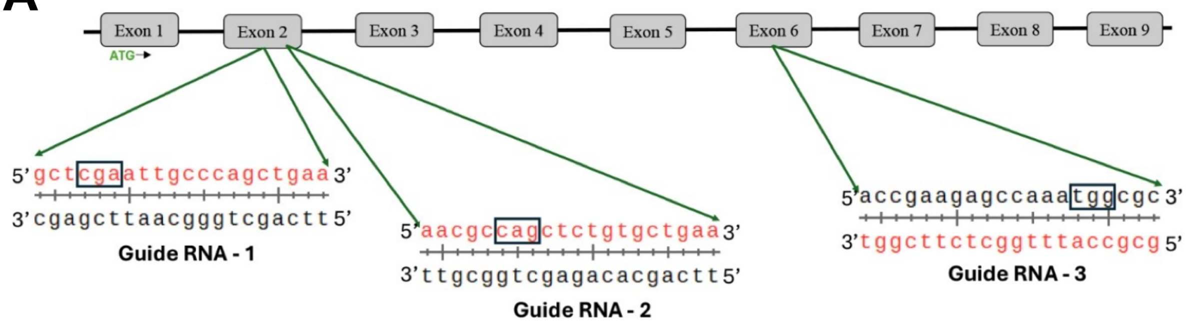


B

| Guide RNA tested | No. of oocytes | No. of blastocysts | Blastocyst percent | KO percent |
|---|---|---|---|---|
| Wild type | 30 | 19 | 63.33 | - |
| Gude #1 | 30 | 20 | 66.67 | 0 % |
| Gude #2 | 30 | 18 | 60.00 | 0 % |
| Gude #3 | 30 | 20 | 66.67 | 80 % |

C

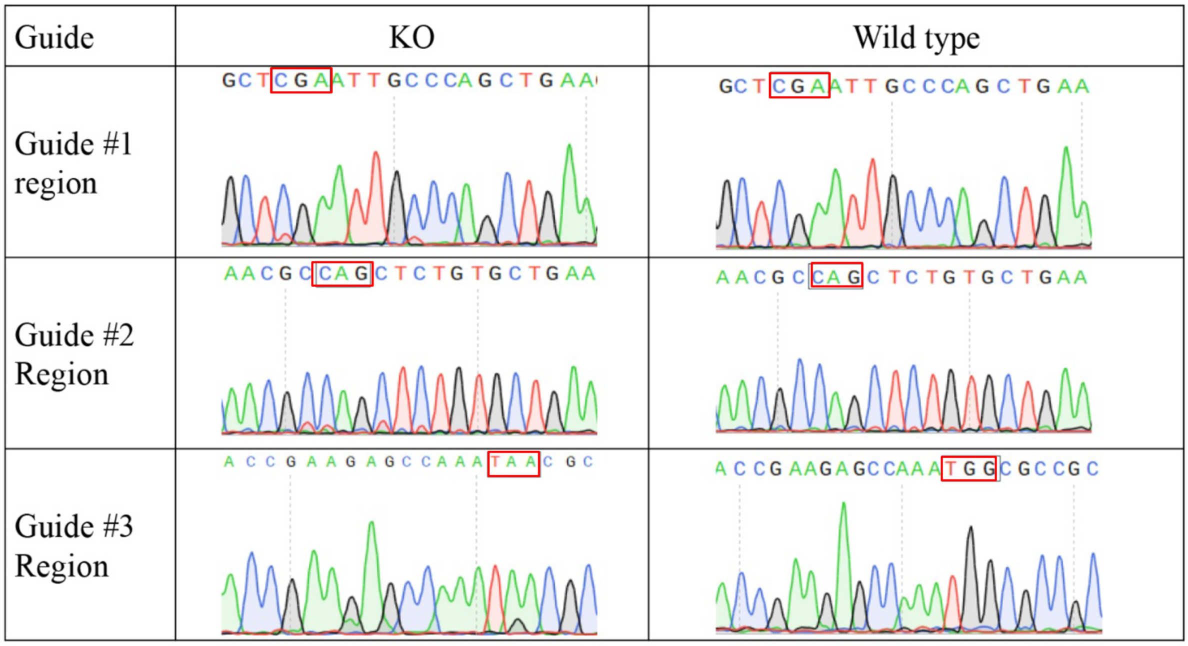


**Figure S1.** Base editor sgRNA design, embryo development outcomes, and sequence validation for targeting porcine *PAX4*. (A) Targeting map of three CRISPR gRNAs across the porcine *PAX4* locus (B) CRISPR guide selection: Blastocyst yield and knockout efficiency of porcine zygotes (C) Sequence verification of CRISPR edits at three gRNA sites in the porcine *PAX4* locus.

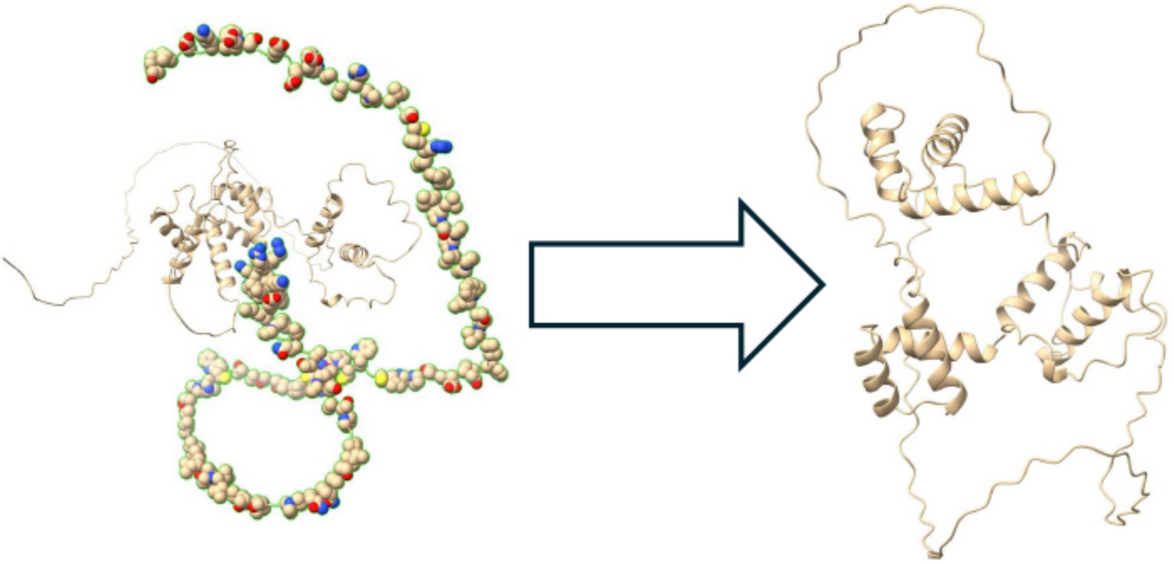

**Figure S2. Predicted 3D structures of wild-type and mutant porcine PAX4 proteins.** The structure of *PAX4* wild-type (**Left**) shows a well-folded region comprising several α-helices, including a portion of the homeobox domain responsible for DNA binding. This domain is highlighted using space-filling (spherical) representation to emphasize the region that is expected to be structurally and functionally critical. In contrast, the *PAX4* mutant (**Right**), generated by introducing a premature stop codon, shows a loss of the homeobox domain and associated structural elements. The absence of this region is expected to result in the loss of the DNA-binding ability of the protein and compromise its role in transcriptional regulation.

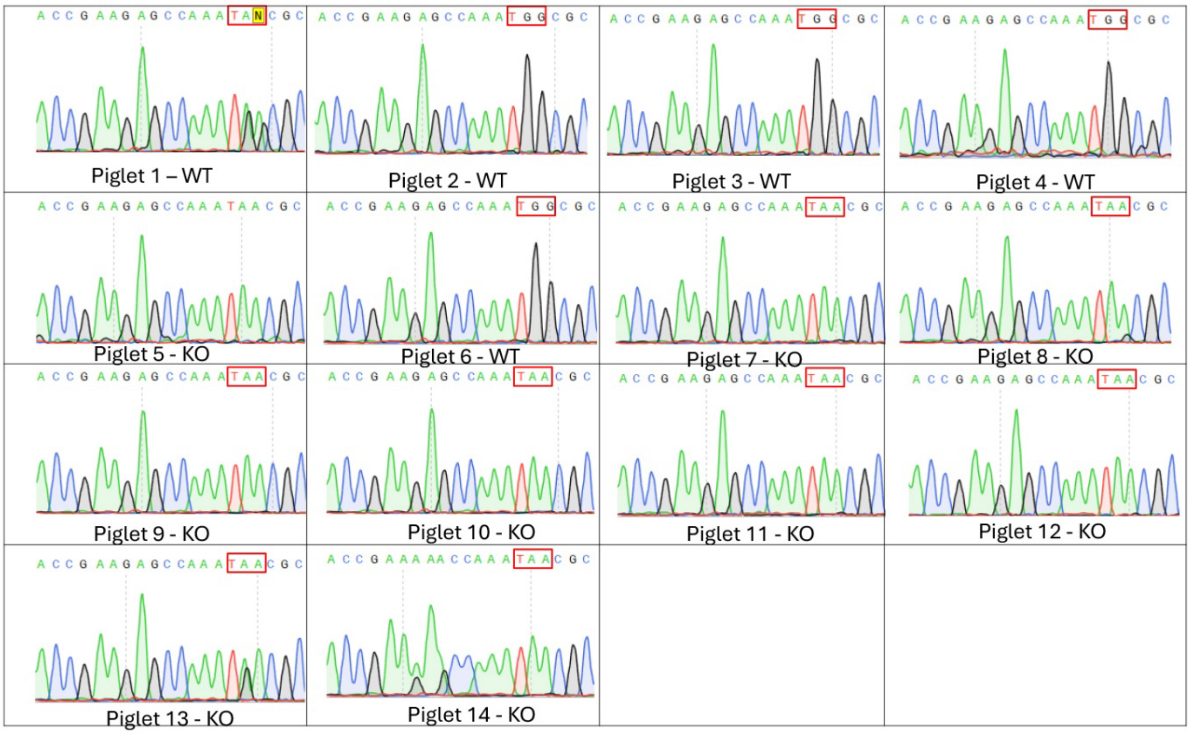


**Figure S3.** Sanger sequencing of PAX4 alleles in WT and BE4 edited piglets**.**

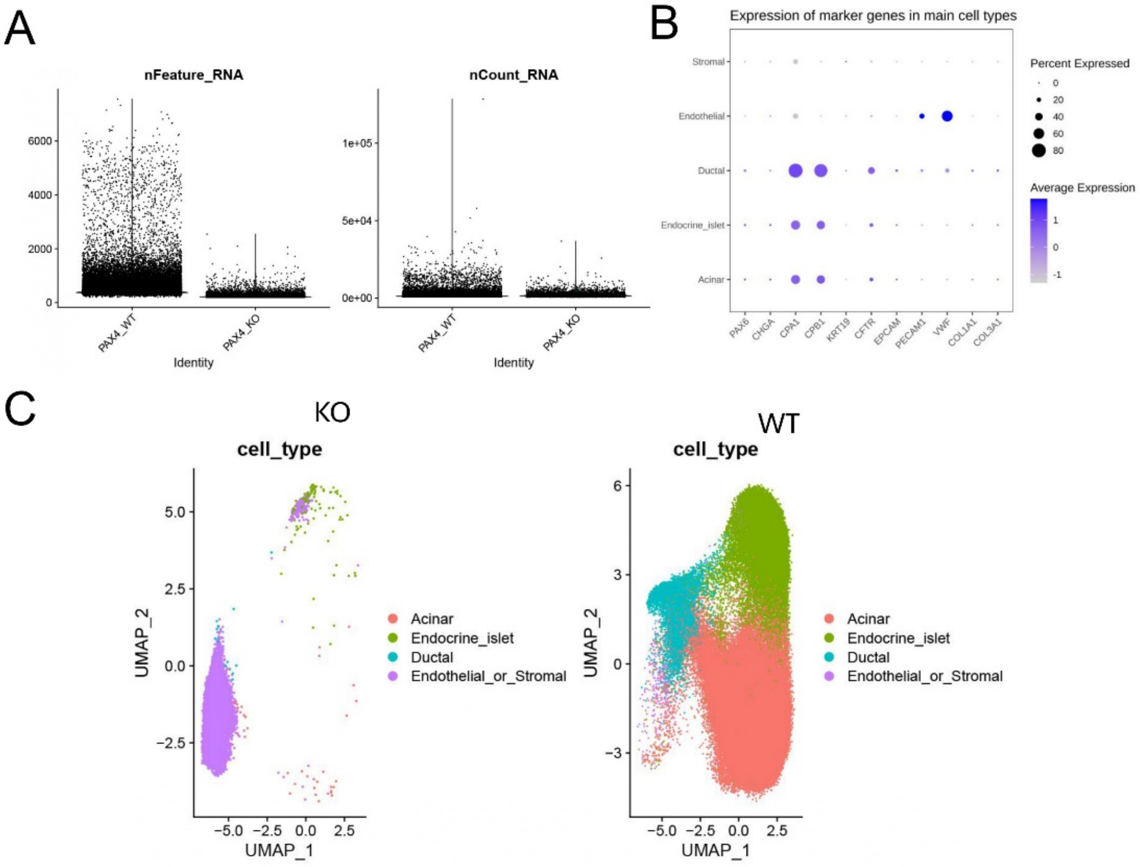


**Figure S4.** SnRNA-seq reveals endocrine-cell depletion and broad transcriptional disruption in *PAX4*-KO neonatal pancreas.
(A) QC metrics. (B) Cell-type marker validation. (C) UMAP by genotype and cell type

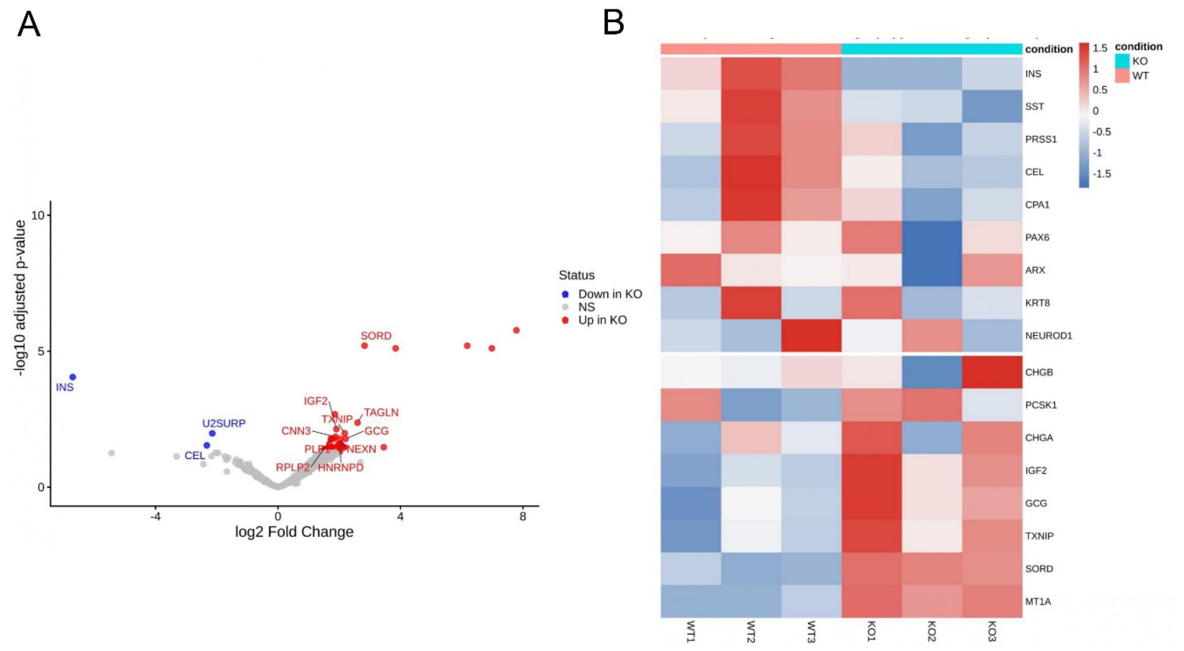


**Figure S5.** Bulk RNA-seq shows loss of endocrine-cell markers and activation of diabetes-associated transcriptional signatures in PAX4-KO neonatal pancreas.

(A) Volcano plot. (B) Targeted heatmap.

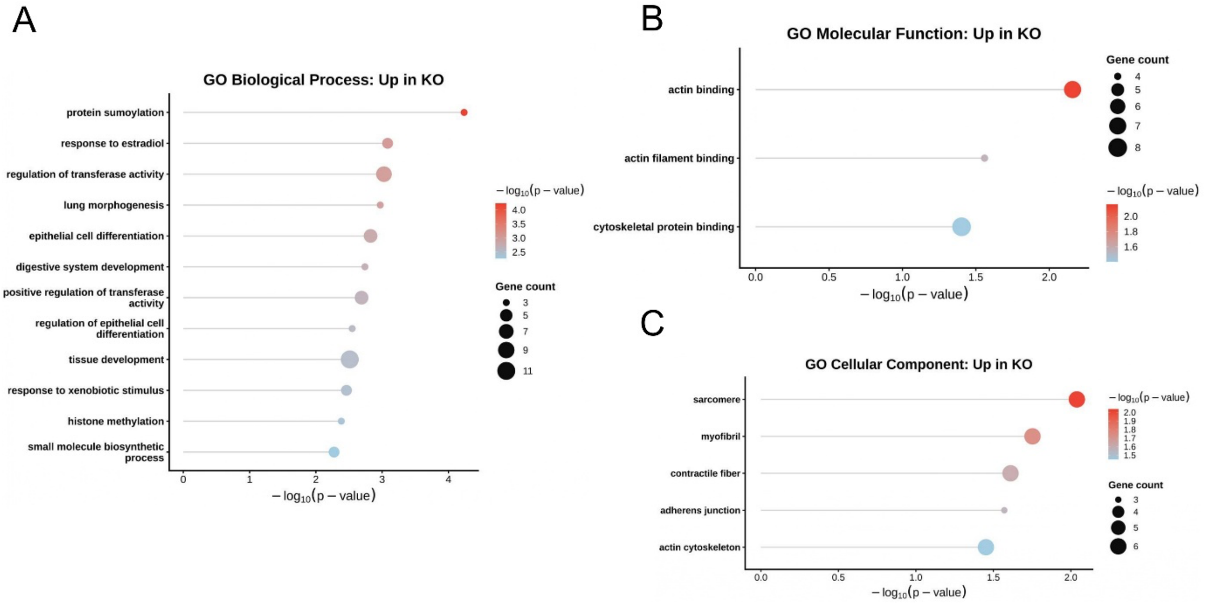


**Figure S6.** Gene ontology enrichment analysis of genes upregulated in PAX4-KO neonatal pancreas.

(A) GO Biological Process enrichment (B) GO Molecular Function enrichment (C) GO Cellular Component enrichment

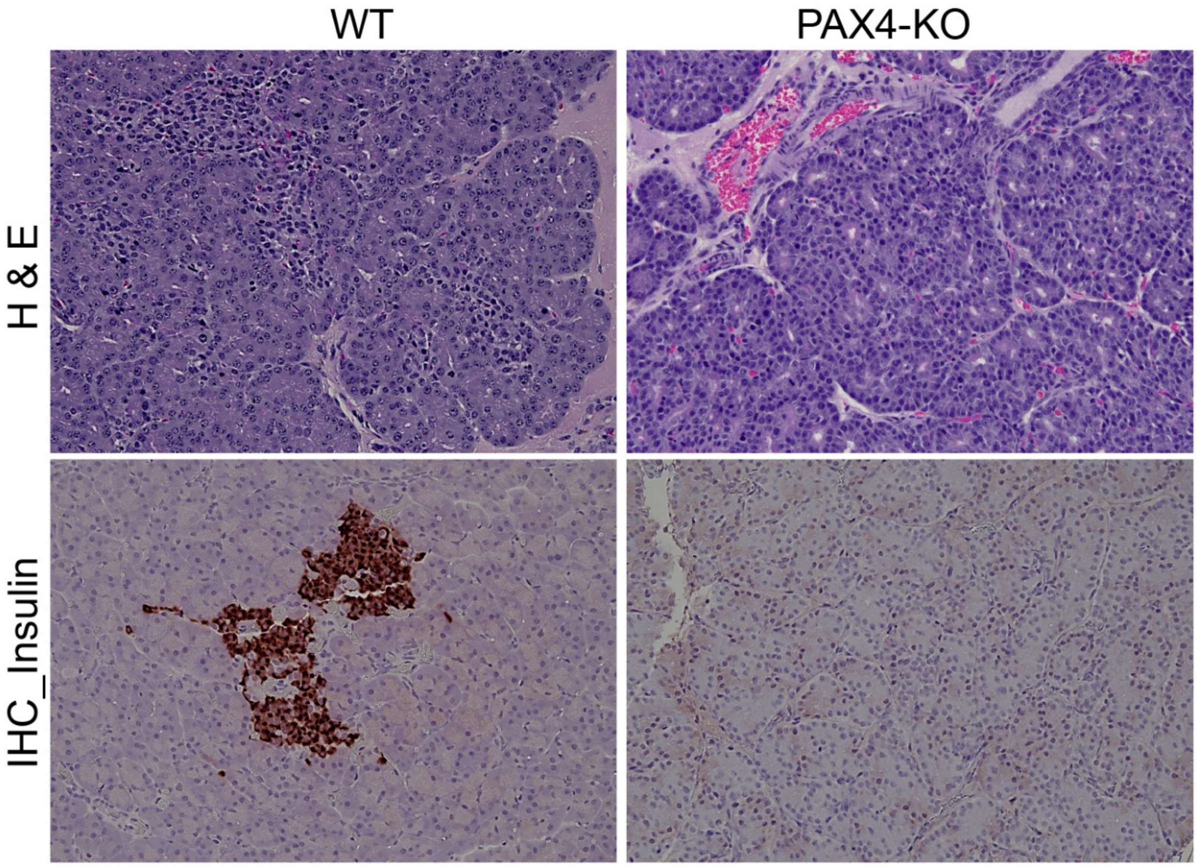


**Figure S7.** Representative H&E histology and insulin immunohistochemistry of WT and *PAX4*-KO neonatal pancreas showing loss of insulin-positive islets in *PAX4*-KO

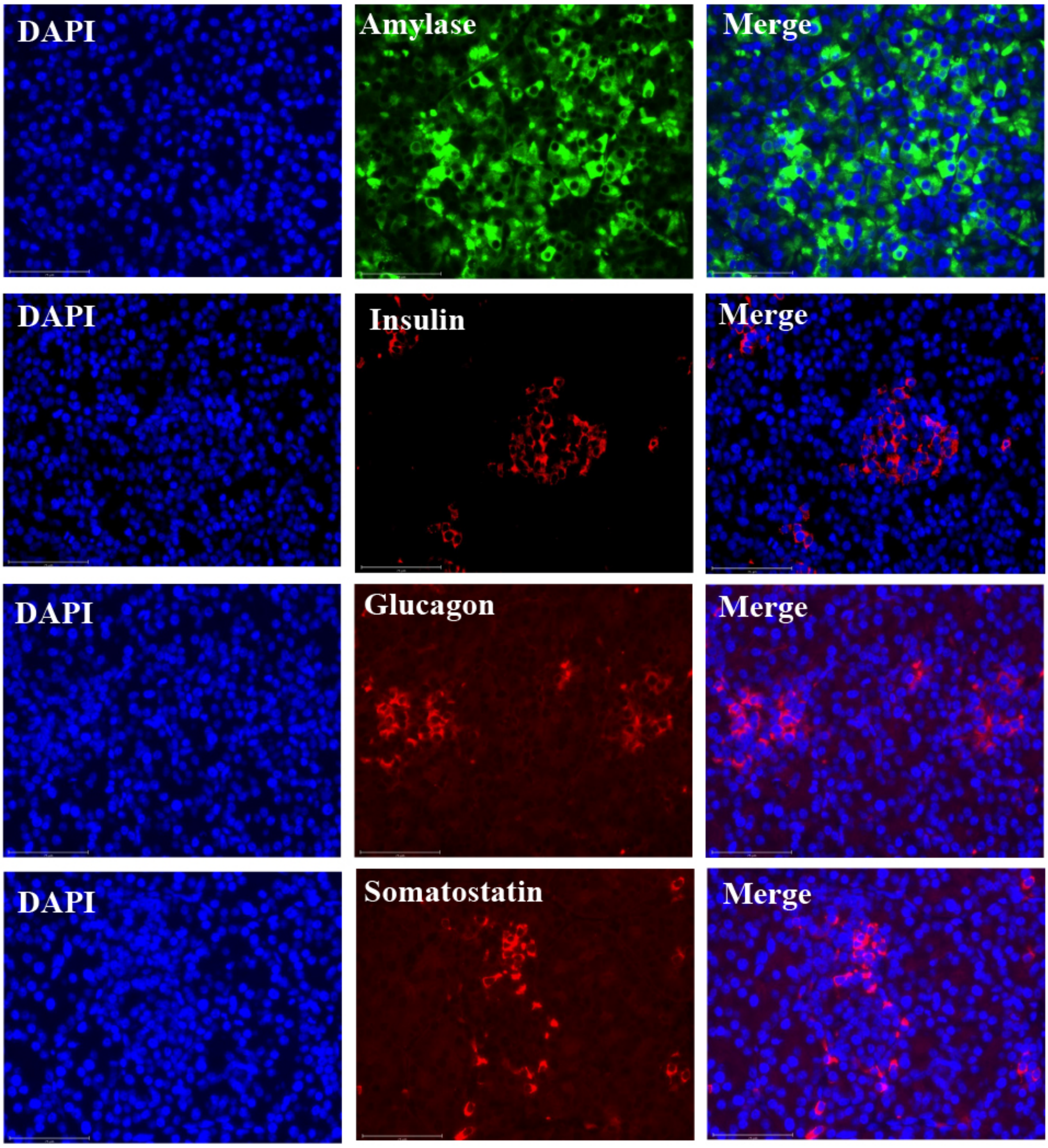


**Figure S8**. Immunofluorescence staining of WT neonatal pancreas showing exocrine amylase- and endocrine hormone-positive cells (insulin, glucagon, somatostatin; 40X)

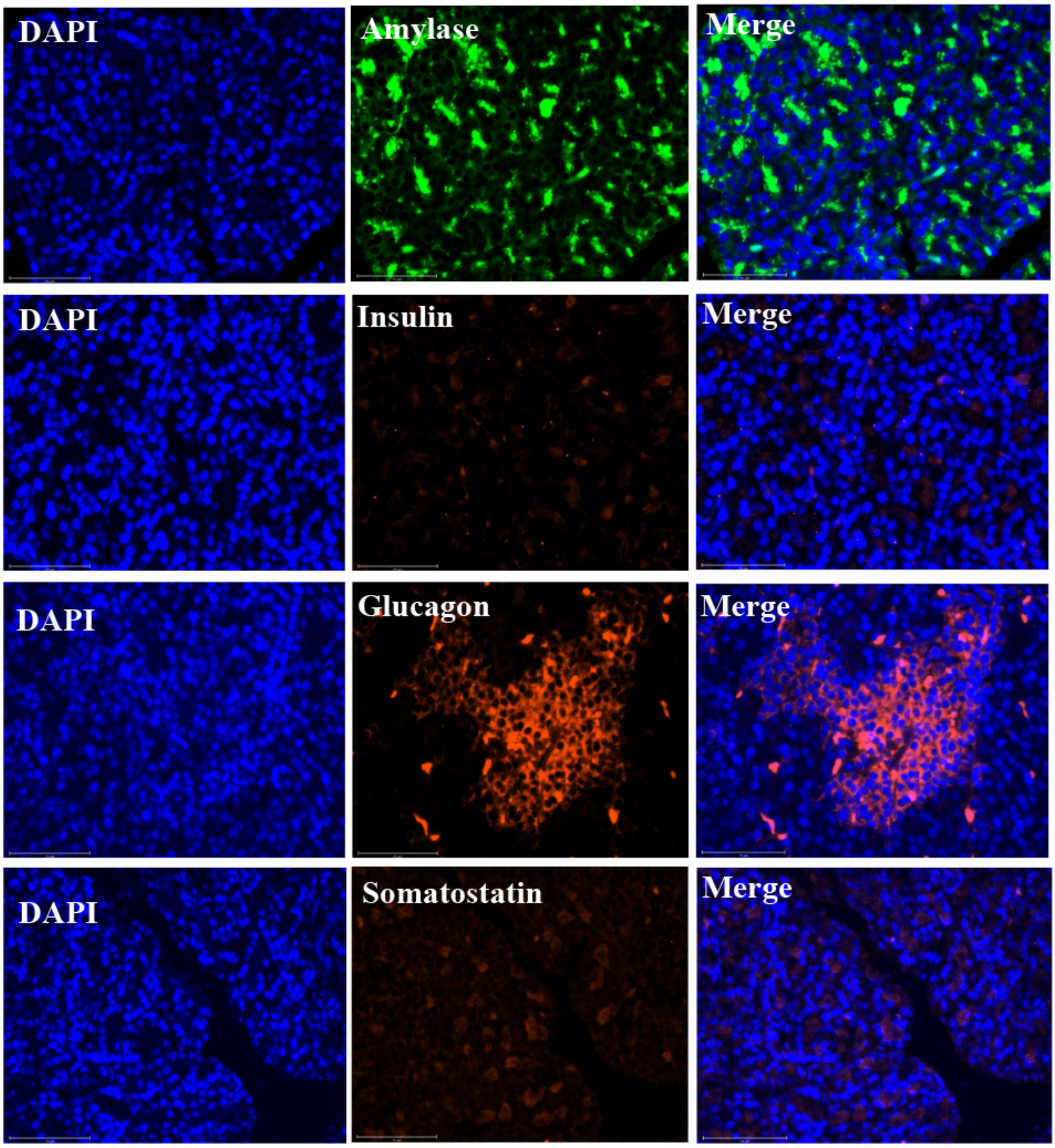


**Figure S9.** Immunofluorescence of *PAX4*-KO pancreas showing preserved amylase staining with loss of insulin- and somatostatin-positive endocrine cells (40X).

Note: The secondary antibodies used for glucagon and somatostatin in Supplementary figures 5 and 6 are different from those used in the main text: insulin was detected with donkey anti-rabbit (Life Technologies, A10042), and somatostatin with donkey anti-goat (Life Technologies, A11057).

**Table S1.** Overview of Guide RNA Sequences and Screening Primers

| | Guide RNA/Primer | Sequence (5′-3′) |
|---|---|---|
| | Guide RNA #1 | GCTCGAATTGCCCAGCTGAA |
| | Guide RNA #2 | AACGCCAGCTCTGTGCTGAA |
| Guide RNA #1 and #2 | Outer_Forward | CCTTACCTGTTCCTGCTTCCTA |
| | Outer_Reverse | TGCCAAGTACAAAGGGATCTTC |
| | Inner_Forward | GCAAGATCCTAGGGCGTTACTA |
| | Inner _Reverse | GTGGCACTTACACTGGGAGTCT |
| | Guide RNA #3 | GCGCCATTTGGCTCTTCGGT |
| Guide RNA #3 | Outer_Forward | AGCAGAAATGAGACAAGAGCCT |
| | Outer_Reverse | TGTGTAGCCCAGCATTTAACAC |
| | Inner_Forward | CAGGTCCGGTGAGAACAGTAG |
| | Inner _Reverse | AGAGGATAGGTGAGTGGAGTGG |

**Table S2.** Antibody Panel and Working Dilutions

| Primary Antibody | | | | |
|---|---|---|---|---|
| S. No. | Antibody | Host | Dilution tested | Catalogue detalis |
| 1 | Anti_insulin | Guinea pig | 1:150 | DAKO (A0564) |
| 2 | Anti_glucagon | Rabbit | 1:200 | Sigma Aldrich (ZRB1115) |
| 3 | Anti_Somatostatin | Rabbit | 1:300 | DAKO (A0566) |
| 4 | Anti_Amylase | Goat | 1:150 | Santa Cruz Biotech (SC-12821) |
| Secondary Antibody | | | | |
| S. No. | Antibody | Species | Catalogue details | |
| 1 | Anti_insulin | Donkey anti_guinea pig | Jackson immunoResearch (706-165-148) | |
| 2 | Anti_glucagon | Goat anti_mouse | Life Technologies (A11029) | |
| 3 | Anti_somatostatin | Donkey anti_rabbit | Novus Biologicals (NBP 1-75638) | |
| 4 | Anti_Amylase | Donkey anti_goat | Novus Biologicals (NBP 1-74824) | |

**Table S3.** Statistical analysis plan and outputs for phenotypic, survival, and serum biochemistry data

| S. No. | Parameter | Normal Distribution (Shapiro wilk's test) | Homoscedasticity (Lavene's test) | Statistical analysis | Statistical output |
|---|---|---|---|---|---|
| 1 | Body weight | Yes (P>0.05) | Yes (P>0.05) | ANCOVA | B/w groups<br>Genotype - P<0.001<br>Timepoint - P=0.176<br>Interaction - P<0.05<br>Within the group<br>WT (Timepoints) - P<0.01<br>KO (Timepoints) - P<0.01 |
| 2 | Body Condition Score (BCS) | No (P<0.05) | No (P<0.05) | Aligned Rank Transform method | Genotype - P<0.001<br>Timepoint - P<0.001<br>Interaction - P=0.08 |
| 3 | Blood Sugar (On-farm testing) | Yes (P>0.05) | Yes (P>0.05) | ANCOVA | Genotype - P<0.001<br>Timepoint - P<0.001<br>Interaction - P<0.05 |
| 4 | Survival curve | - | - | Mantel–Cox test | Genotype - P = 0.033 (<0.05) |
| 5 | Serum components | Unequal and small groups | | Kruskal-Wallis test | - |
